# A common pattern of influenza A virus single cell gene expression heterogeneity governs the innate antiviral response to infection

**DOI:** 10.1101/858373

**Authors:** J. Cristobal Vera, Jiayi Sun, Yen Ting Lin, Jenny Drnevich, Ruian Ke, Christopher B. Brooke

**Affiliations:** Department of Microbiology, University of Illinois at Urbana-Champaign, Urbana, IL; Carl R. Woese Institute for Genomic Biology, University of Illinois at Urbana-Champaign, Urbana, IL; T-6, Theoretical Biology and Biophysics, Los Alamos National Laboratory, Los Alamos, NM; High-Performance Biological Computing at the Roy J. Carver Biotechnology Center, University of Illinois at Urbana-Champaign, Urbana, IL

## Abstract

Viral infection outcomes are governed by the complex and dynamic interplay between the infecting virus population and the host response. It is increasingly clear that both viral and host cell populations are highly heterogeneous, but little is known about how this heterogeneity influences infection dynamics or viral pathogenicity. To dissect the interactions between influenza A virus (IAV) and host cell heterogeneity, we examined the combined host and viral transcriptomes of thousands of individual, single virion-infected cells. We observed complex patterns of viral gene expression and the existence of multiple distinct host transcriptional responses to infection at the single cell level. Our analyses reveal that viral NS segment gene expression diverges from that of the rest of the viral genome within a subset of infected cells, and that this unique pattern of NS segment expression can play a dominant role in shaping the host cell response to infection. Finally, we show that seasonal human H1N1 and H3N2 strains differ significantly in patterns of host anti-viral gene transcriptional heterogeneity at the single cell level. Altogether, these data reveal a common pattern of viral gene expression heterogeneity across human IAV subtypes that can serve as a major determinant of antiviral gene activation.

## INTRODUCTION

RNA virus populations typically contain an enormous amount of sequence diversity due to an absence of virally encoded proofreading activity (1). In some cases, this genetic diversity can significantly influence infection outcomes (2–4). In addition, influenza A viruses (IAV) also exhibit substantial heterogeneity in the gene expression patterns of individual virions. Most IAV virions are only capable of expressing variable, incomplete subsets of viral genes (5,6). The production and gene expression patterns of these semi-infectious particles (SIPs) can vary significantly between IAV strains (7). Altogether, the high degree of diversity within IAV populations means that patterns of viral gene expression can vary significantly between individual infected cells.

Numerous studies have leveraged recent advances in single cell analysis methods to assess the extent of cellular heterogeneity present during infection by a variety of viruses, including IAV (8–17). These studies connect to a larger body of work that has begun to explore single cell heterogeneity within different host cell populations and tissues (18–20). To date, efforts to characterize single cell heterogeneity during IAV infection have been hampered by limitations in the number of cells analyzed or experimental design choices that complicated analysis. The extent to which IAV population diversity influences patterns of single cell heterogeneity and the ways in which these patterns shape broader infection dynamics and outcomes remain poorly understood.

Here, we query the combined viral and host transcriptomes from thousands of individual cells, each infected by a single virion. This high-resolution dissection of viral and host gene expression patterns reveals that the transcriptional responses of individual infected cells can be highly divergent, resulting from the interplay between underlying cellular and viral heterogeneity. Thus, the host response to IAV infection consists of a heterogenous assemblage of highly variable single cell responses. This approach also reveals critical differences in the interactions of H1N1 and H3N2 viruses with the host antiviral machinery at the single cell level and implicates variability in NS segment expression as a major driver of the innate antiviral response to IAV infection. Altogether, these results establish the interplay between viral and host cell heterogeneity as a critical determinant of cellular infection outcomes.

## RESULTS

### Generation of viral and host transcriptional data from thousands of singly infected cells

To assess the effects of viral population heterogeneity on the host response to infection, we examined the combined viral and host transcriptional profiles from thousands of single infected cells. To focus on the effects of viral heterogeneity, we wanted to remove the variability that could arise from variation in cellular MOI (21). To ensure that >99.99% of infected cells were each infected with a single virion, we infected A549 cells with the 2009 H1N1 pandemic strain A/California/07/2009 (Cal07) at an MOI of 0.01, and blocked secondary spread in the culture through the addition of NH4Cl (22). This resulted in only a tiny fraction of cells being infected, so to obtain sufficient numbers of infected cells for our analysis, we enriched infected cells by sorting based on surface expression of HA and/or M2 (**Fig. 1A**). This approach had the added benefit of allowing us to sort HA-M2-bystander cells from the same culture to assess the paracrine effects of infection. In parallel, we sorted a group of mock-infected cells as a control.

**Fig. 1.**
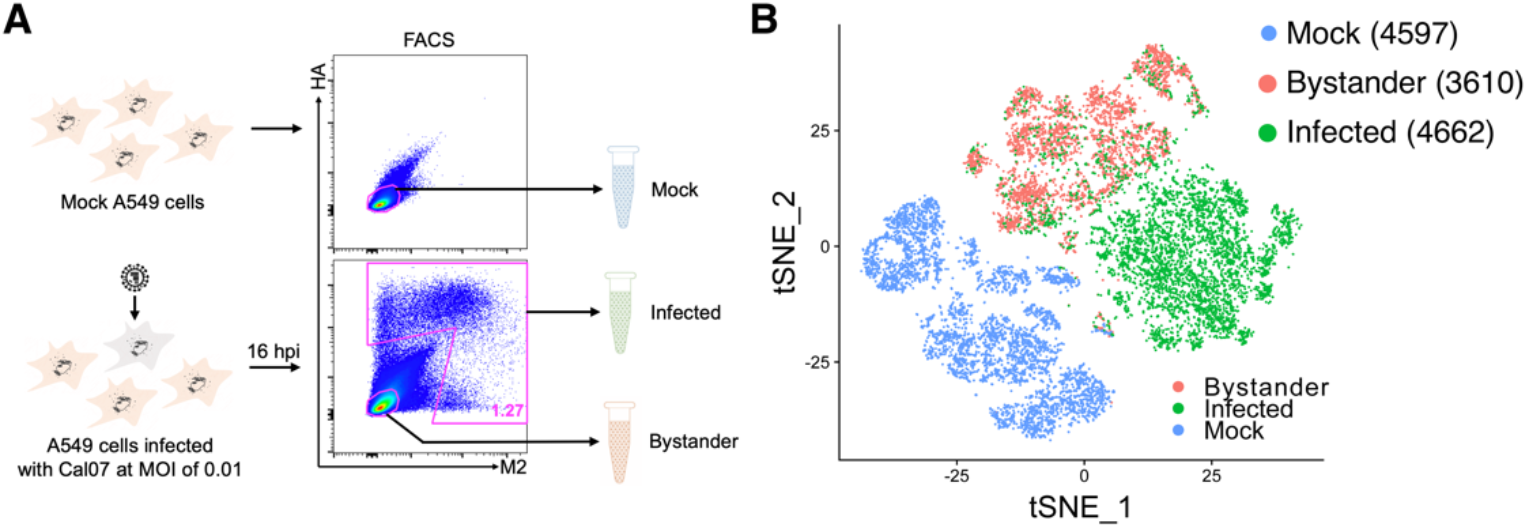
Generation of viral and host transcriptional data for thousands of singly infected cells. **(A)** Schematic depicting our strategy for generating single cell RNAseq libraries from thousands of cells infected at low MOI. In brief, we infect A549 cells with Cal07 at MOI=0.01 to ensure that infected cells are infected with a single virion. We then block secondary spread with NH4CI treatment to make sure infection timing is uniform across all infected cells. Finally, we sort “infected” and “bystander” cells based on surface expression of HA and/or M2 and immediately generate single cell RNAseq libraries from these sorted cell populations using the 10X Chromium device. In parallel, mock cells are sorted and used as uninfected controls. **(B)** tSNE dimensionality reduction plot showing the extent of overlap of the indicated cell populations clustered based on transcriptional similarity.

We used the 10X Genomics Chromium platform to generate oligo-dT-primed single cell RNAseq libraries for mock, infected (HA+ and/or M2+), and bystander (HA-M2-) cells and sequenced them on an Illumina Novaseq. We demultiplexed the reads and mapped them to a customized hybrid reference containing both human and influenza sequences/annotation. Following quality control filtering, we had high quality combined viral and host transcriptomes from 4662 single virion-infected cells, as well as 3610 bystander cells, and 4597 mock cells. Clustering of these three libraries by transcriptional similarity revealed that mock, bystander, and infected cells largely clustered independently, as would be expected, though a subset of infected cells grouped closely with the bystander cells (**Fig. 1B**).

### Enormous single cell heterogeneity in viral gene expression patterns

We first asked whether we observed the same degree of heterogeneity in viral gene expression between infected cells that has been reported previously (6,10,12). We calculated the fraction of all transcripts within each cell that were viral in origin (**Fig. 2A**). Similar to prior studies, total viral transcript levels ranged enormously between individual cells: from <1% up to ~90%. Notably, this heterogeneity arose under conditions where both viral input (1 virion/cell) and infection timing were equivalent across all cells.

**Fig. 2.**
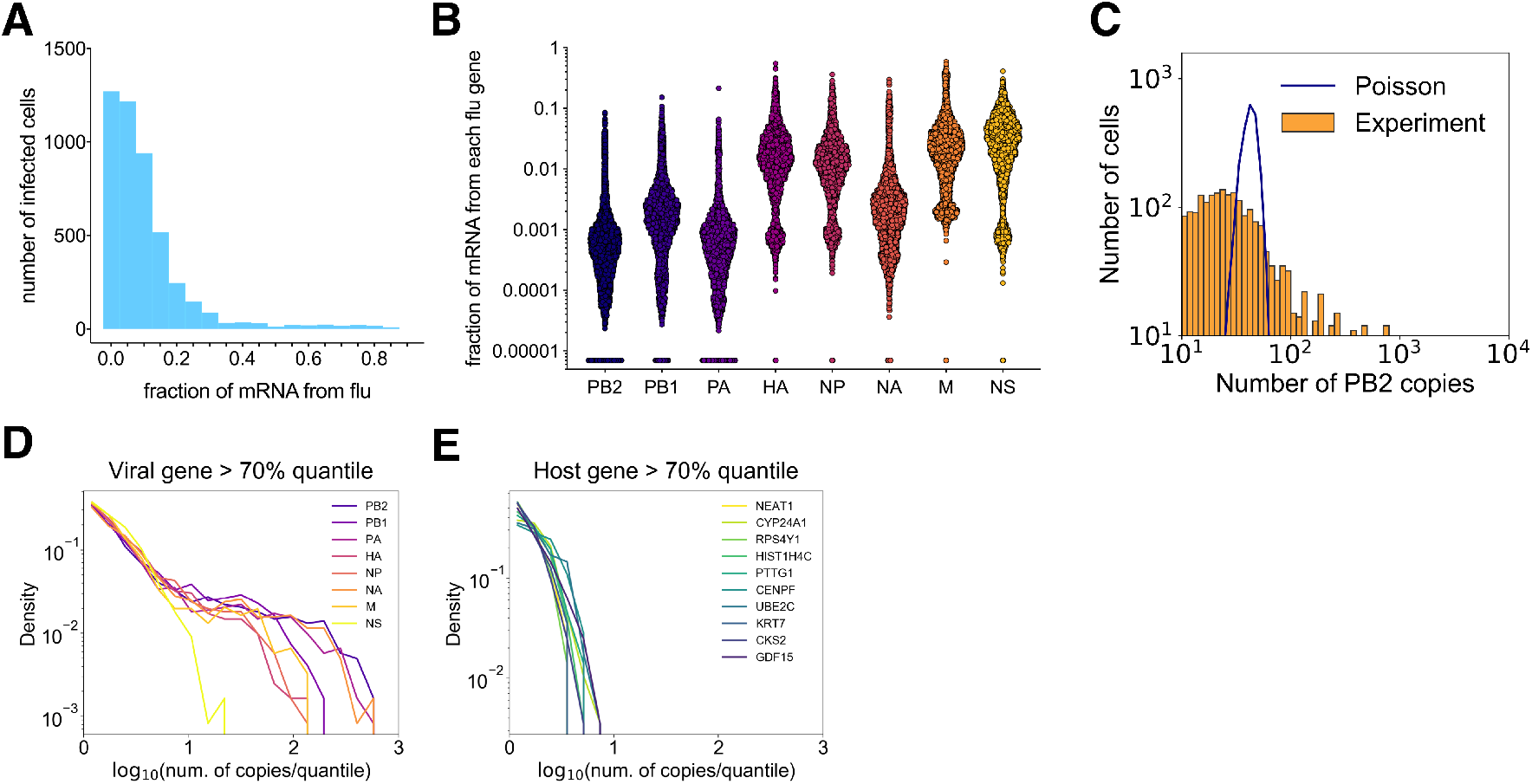
Enormous heterogeneity in viral gene expression patterns. **(A)** Distribution of Cal07-infected A549 cells, binned by the fraction of total cellular poly(A) RNA that is viral in origin. **(B)** Fraction of total poly(A) RNA per cell that maps to the indicated viral gene segment. Each dot represents a single cell, cells with no detectable reads mapping to the indicated segment are arbitrarily assigned a value of 0.000007 to show up on the log10 scale. **(C)** Distribution of normalized PB2-derived poly(A) RNA counts per cell (orange) compared with a Poisson distribution of equal mean (blue line) on a log-log scale. **(D)** The viral gene expression distributions normalized by the 70^th^ quantile on a log-log scale. **(E)** Same analysis as in (D) but showing the top 10 most abundant host transcripts.

This pattern of extreme cell-to-cell heterogeneity was also observed for individual viral gene segments, again consistent with previous reports (**Fig. 2B**)(12). Due to the short reads obtained using our single cell sequencing approach, we were not able to reliably differentiate between the different viral transcripts generated from individual gene segments (*i.e*. NS1 vs. NEP or M1 vs. M2) so for all analyses we identified all viral transcripts by gene segment of origin. Expression distributions for individual viral gene segments were highly over-dispersed compared with a Poisson distribution (**Fig 2C**). The tails of the viral gene expression distributions were orders of magnitude longer than those of the top 10 most abundant host transcripts (**Fig. 2D,E**), emphasizing that viral gene expression is substantially noisier than host gene expression.

### Significant heterogeneity in the host transcriptional response to infection

We next asked whether the observed variation in viral gene expression levels was associated with variation in host cell transcription. To test this, we first clustered all infected cells based on host transcription patterns. We identified multiple distinct transcriptional response groups to infection by Cal07, suggesting that there is not one standard transcriptional response to IAV infection and that a single cell type can simultaneously generate multiple distinct responses to the same virus population (**Fig. 3A,B**). It should be noted that these cluster definitions are not absolute and that substantial heterogeneity in the normalized expression levels of individual host genes existed within individual clusters.

**Fig. 3.**
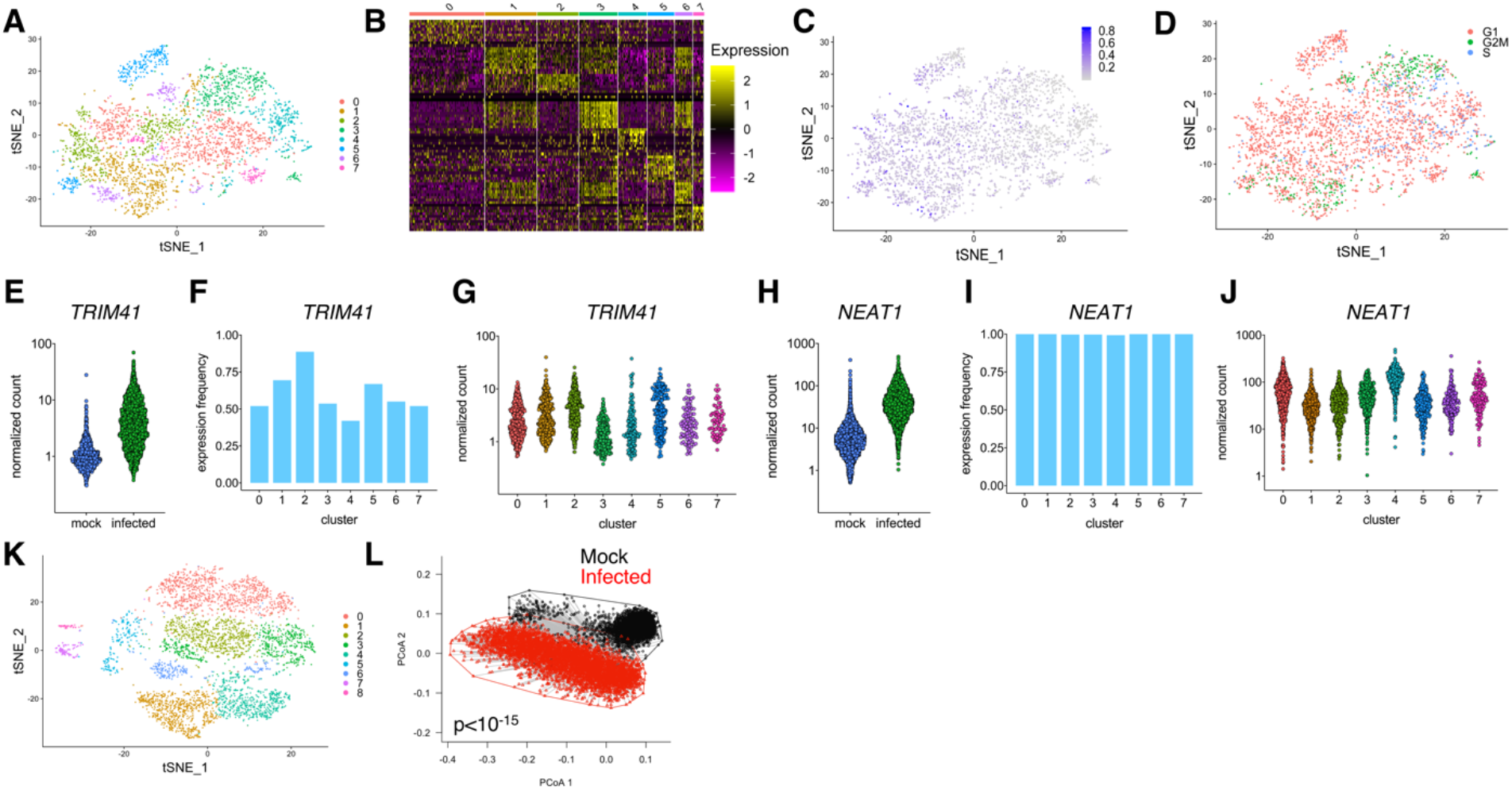
Significant heterogeneity in the host transcriptional response to infection. **(A)** tSNE dimensionality reduction plot showing all Cal07-infected A549 cells clustered based on similarity of host transcriptional patterns. **(B)** Heat map showing differential expression of the top 10 characteristic host genes for each cluster from (A). Individual cells are each represented by a column, grouped by cluster, with individual rows representing relative expression of the top 10 specific host transcripts most significantly associated with each cluster. **(C)** Same tSNE plot of Cal07-infected cells shown in (A) with each cell colored by the proportion of total cellular poly(A) RNA that is viral in origin. **(D)** Same tSNE plot of Cal07-infected cells shown in (A) with each cell colored by predicted cell cycle stage, as determined by the scran package. **(E)** Comparison of normalized per cell counts of TRIM41 between mock and infected cells. **(F)** Fraction of cells in each infected cell cluster from (A) with detectable levels of TRIM41. **(G)** Comparison of normalized per cell counts of TRIM41 between clusters of infected cells shown in (A). **(H)** Comparison of normalized per cell counts of NEAT1 between mock and infected cells. **(F)** Fraction of cells in each infected cell cluster from (A) with detectable levels of NEAT1. **(G)** Comparison of normalized per cell counts of NEAT1 between clusters of infected cells shown in (A). **(K)** tSNE dimensionality reduction plot showing mock A549 cells clustered based on similarity of host transcriptional patterns. **(L)** Principle coordinate axes (PCoA) plot comparing the multivariate dispersions for mock (black) and Cal07-infected (red) A549 cells. The first two axes (PCoA 1 and PCoA 2) in the multivariate homogeneity of group dispersions analysis are used.

We did not observe any clear associations between clustering pattern and total viral transcript levels (**Fig. 3C**); however, the clustering pattern did appear to be partially influenced by cell cycle status (**Fig. 3D**). Cells in G2M phase were primarily found within a distinct subset of clusters (1, 3, and 6) while cells in G1 and S phases exhibited no clear clustering patterns. Consistent with this, the expression levels of a number of host cell cycle-associated genes (*e.g*. CDK1, CENPF, MKI67, PTTG1, and CKS2) were significantly elevated in those same clusters (**Table S1**). While it is not surprising that cell cycle status would contribute to transcriptional heterogeneity during infection, these data highlight how little is known about how cell cycle status may influence the cellular response to infection.

Most of the observed heterogeneity could not be simply explained by cell cycle, however. The induction of numerous host genes known or likely to be involved in shaping IAV infection outcome varied significantly between clusters. One notable example is TRIM41, a protein previously shown to inhibit IAV replication by marking viral NP for degradation (23). TRIM41 is clearly upregulated in response to infection at the bulk level (**Fig. 3E**). At the single cell level, both the frequencies and the overall expression levels of this transcript differed significantly between infected cell clusters (**Fig. 3F,G**). We also observed substantial variation between clusters in upregulation of the long non-coding RNA NEAT1, which is involved in inflammasome formation, regulation of cytokine and chemokine expression, and nuclear paraspeckle formation (24–29). NEAT1 expression was upregulated during infection at the bulk level (**Fig. 3H**), but levels were significantly higher in clusters 0 and 4 compared with other infected cells (adjusted p values = 2.01*10^-12^ and 2.91*10^-112^ respectively) (**Fig. 3I,J**). Thousands of other host genes exhibited similarly heterogeneous patterns of expression between individual infected cells, illustrating how key drivers of infection outcomes may be primarily expressed by limited subsets of infected cells.

We also observed the existence of several distinct clusters within mock cells (**Fig. 3K**), leading us to ask whether the heterogeneity we observed in infected cells was simply a reflection of the intrinsic heterogeneity of the cell population prior to infection. In other words, does infection actually increase the overall heterogeneity in host gene expression patterns beyond that seen in mock cells? To quantify and compare heterogeneity in overall host transcription between the two cell populations, we calculated the multivariate homogeneity of dispersions for mock and infected cells using expression of all host genes (**Fig. 3L**)(30). We found that infection significantly increased the overall single cell heterogeneity in host cell transcription patterns (p<10^-15^ by pairwise t test). Altogether, our data are consistent with a model in which the interaction between viral population heterogeneity and pre-existing host cell heterogeneity gives rise to multiple distinct transcriptional responses at the single cell level.

### Many infected cells have undetectable levels of one or more viral transcripts

The vast majority of IAV virions fail to express one or more viral genes, resulting in the expression of variable, incomplete subsets of viral gene products under low MOI conditions (5–7,31). We asked whether this variation in functional viral gene content within individual cells contributes to the observed heterogeneity in host gene transcription. To assess the presence or absence of each individual viral gene within infected cells while avoiding false positives due to RNA cross-contamination, we took advantage of the fact that the vast majority of “bystander” cells in our analysis were not infected but were collected from the same culture vessel as our infected cells. We used these “true negative” cells to empirically determine the distribution of viral read counts that could arise from mRNA cross-contamination during infection and drop-making (12). We then used these values to generate cutoff thresholds that could be used to differentiate true positives from false positives.

Using these cutoffs, we found that the fractions of infected cells that failed to express detectable transcripts ranged from ~35% to nearly zero for the individual viral gene segments (**Fig. 4A**). It is highly likely that our method of enriching for infected cells by sorting based on high level HA and/or M2 expression significantly biased these results. Regardless, we still observed substantial numbers of infected cells that fail to express detectable levels of individual viral transcripts, with roughly half of all infected cells failing to express detectable levels of at least one viral gene (**Fig. 4B**). We were surprised to see so many cells that lacked detectable levels of the polymerase transcripts because we expected that sorting infected cells based on surface protein expression would bias cell collection against cells that could not synthesize new polymerase complexes. One caveat with this approach is that we assessed transcript levels at 16 hpi and thus could have missed transient early expression of viral genes.

**Fig. 4.**
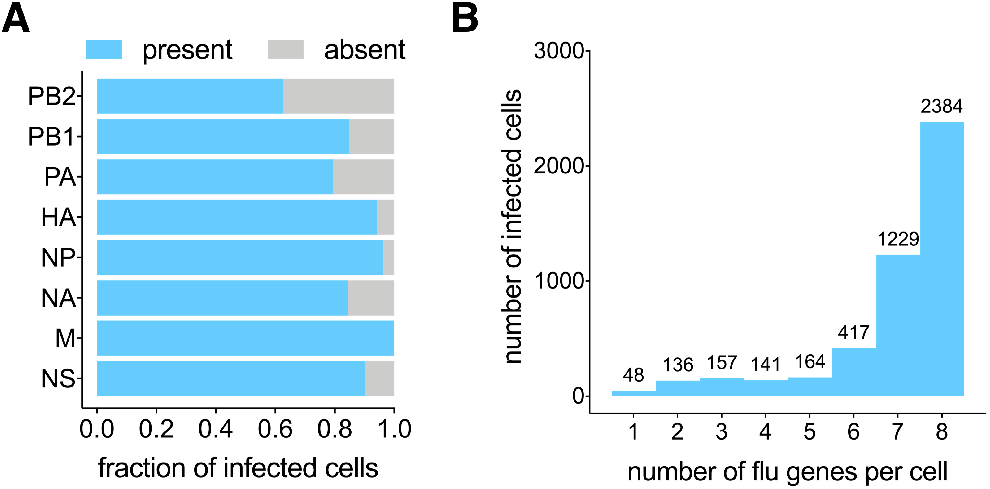
Many infected cells have undetectable levels of one or more viral transcripts. **(A)** Fraction of all Cal07-infected cells that have detectable levels of transcripts derived from the indicated viral gene segments. **(B)** All Cal07-infected cells binned by the total number of detectable viral gene segments, with the actual numbers of cells in each group detailed above.

### Dissecting the effects of individual viral genes on the host transcriptional response to infection

We next asked how the expression of individual viral genes influences the overall host response to infection. We grouped all Cal07-infected cells into positive and negative populations based on the expression of each individual viral gene segment and used two distinct differential single cell gene expression analysis methods (MAST and NBID) to compare host transcript expression between the two infected cell populations (32,33). For each viral gene, we generated a list of host transcripts that were reported as significantly different between positive and negative cells by both NBID and MAST (**Table S2-S9**). This approach allowed us to tease out the effects of individual viral gene segments from the more general effects of infection.

We identified hundreds of host genes with expression levels that varied significantly based on the expression status of individual viral genes (**Fig. 5A**). The one exception was the M segment, most likely due to the lack of sufficient cells lacking M transcript for the analysis. Closer examination revealed that many of these hits were found for multiple viral genes, suggesting that they may correlate with overall viral gene expression levels or co-expression of multiple viral gene segments. When we focused on the host genes that only exhibited differential expression in association with the expression status of a single viral gene segment, we found that most were associated with the PA, NP, or NS segments (**Fig. 5B, Table S10**).

**Fig. 5.**
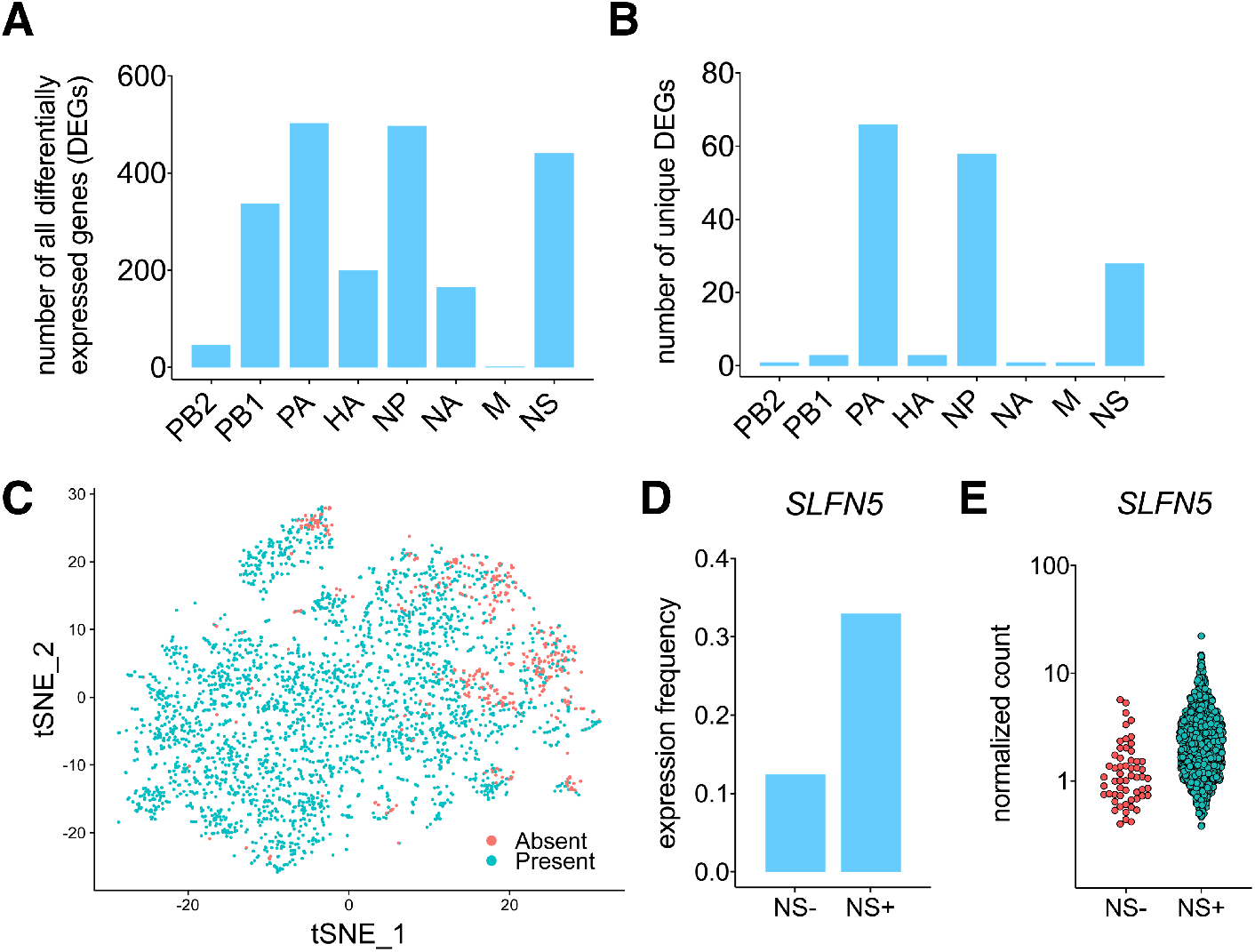
Dissection of the effects of individual viral genes on the host transcriptional response to infection. **(A)** The number of host transcripts for which expression levels significantly differ depending on whether the indicated viral gene segment is present or not, according to both MAST and NBID. **(B)** Same as in (A) but removing host genes that are differentially regulated by the expression status of more than one viral gene segment. **(C)** tSNE plot of all Cal07-infected A549 cells colored based on whether NS segment-derived transcripts are detected (Cyan) or not detected (Salmon). **(D)** Fraction of NS- and NS+ Cal07-infected A549 cells with detectable levels of SLFN5. **(E)** Distribution of SLFN5 normalized counts in cells shown in (D).

We focused on the NS segment, as the NS-encoded NS1 protein plays a well-established, multi-functional role in manipulating the host cell environment and the anti-viral response and could thus serve as a positive control for our approach (34–37). We found that ~10% of infected cells had background or undetectable levels of NS segment transcript (**Fig. 5C**). We observed nearly 450 host genes whose expression was significantly correlated with NS expression status, however only 28 of these were uniquely influenced by NS. Notably, this list included SLFN5 (down in NS-negative infected cells; FDR=3.22*10^-42^), an interferon-stimulated gene (ISG) shown to negatively regulate STAT1-dependent anti-viral gene transcription (38). We observed that both the frequency and normalized per-cell expression levels of SLFN5 were significantly decreased in infected cells that failed to express NS compared with those that did express NS (**Fig. 5D,E**). Expression of specific IFNs or other ISGs was not significantly affected by NS expression status, echoing the previous finding that lack of NS expression does not deterministically result in increased expression of these genes (12,13). These results both validate the utility of our approach and identify the suppressor of antiviral gene transcription SLFN5 as a novel host target of the NS gene segment.

### H1N1 and H3N2 viruses share a common pattern of single cell viral gene expression heterogeneity

Since productively infected cells should express products from all viral gene segments, we asked how variation in viral gene expression within infected cells that express all viral genes affects the host transcriptional response. We examined the degree to which individual pairs of viral genes were correlated with each other (**Fig. 6A, S1**). For Cal07, we found that most pairs of viral transcripts were highly correlated. In particular all pairwise comparisons of the polymerase components exhibited r^2^ values of 0.796 or higher. In contrast, the NS segment was not generally positively correlated with the other segments (**Fig. 6B**).

**Fig. 6.**
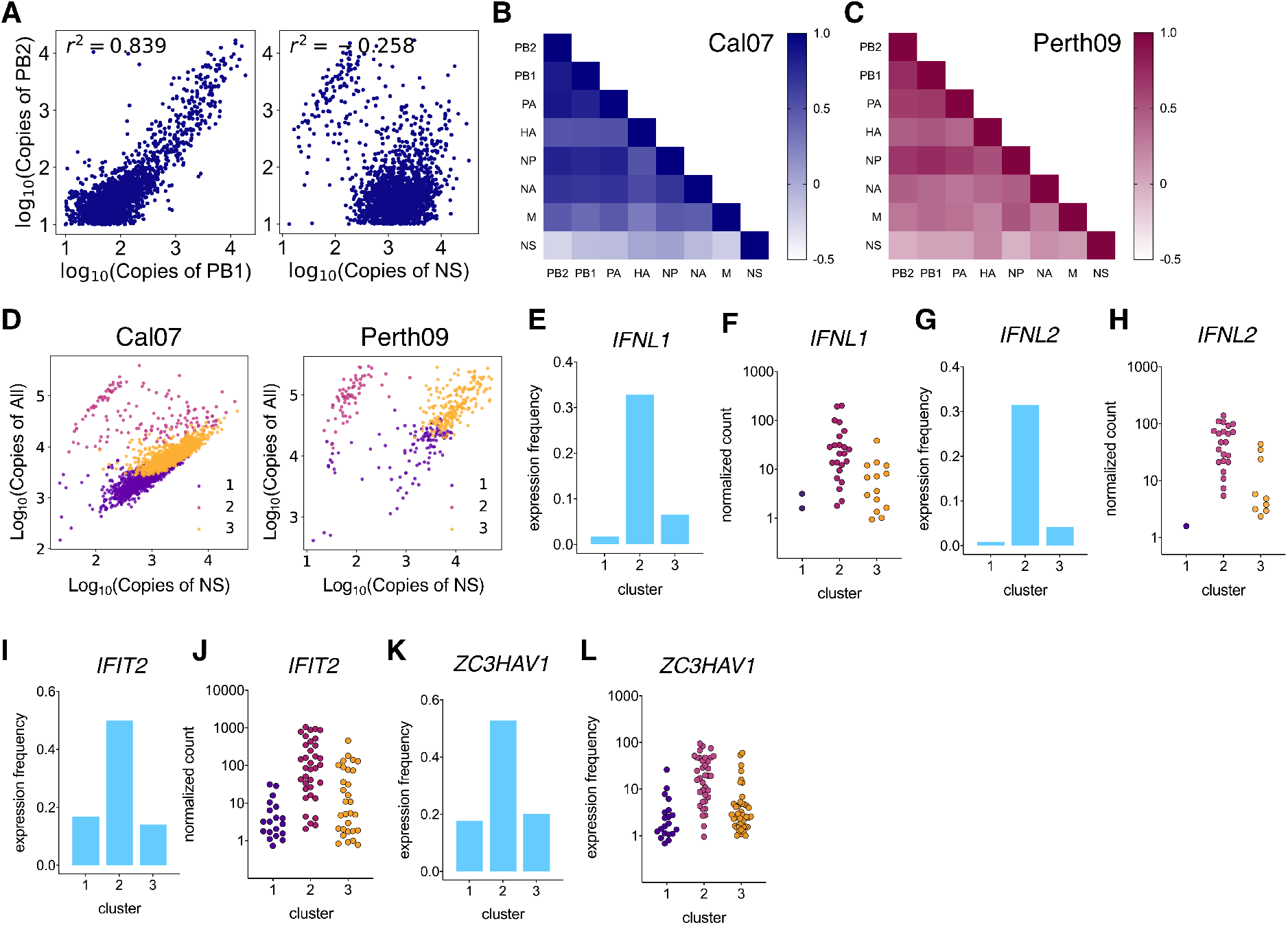
Antiviral gene transcription is governed by a common pattern of single cell viral gene expression heterogeneity. **(A)** Pairwise comparisons between normalized copy numbers of the indicated viral transcripts within Cal07-infected cells in which all viral gene segments are detected. Shown here are the comparisons of PB2 with PB1 (left) and NS (right), along with r^2^ values of correlation. **(B)** Correlation values plotted as heatmap for all pairwise comparisons of viral transcripts within Cal07-infected cells that express all viral genes. **(C)** Same as in (B) for Perth09-infected cells. **(D)** All Cal07- (left) and Perth09- (right) infected A549 cells that are positive for all eight viral gene segments, plotted as a function of total viral gene expression versus NS segment expression. Each dot represents the viral transcriptome of a single cell, colored by cluster identity of K-means clustering based on viral gene expression. Cluster IDs are shown in the legend. **(E)** The fraction of Perth09-infected cells in the clusters shown in (D) with detectable levels of IFNL1. **(F)** Distribution of IFNL1 normalized counts in cells shown in (E). **(G, H)** Same as in (E,F) for IFNL2. **(I,J)** Same as in (E,F) for IFIT2. **(K,L)** Same as in (E,F) for ZC3HAV1.

To determine whether this pattern was a unique feature of the Cal07 strain, we used the same experimental approach detailed above to generate single cell RNAseq libraries for A549 cells infected at low MOI with the human seasonal H3N2 strain A/Perth/16/2009 (Perth09). Analysis of 1197 Perth09-infected cells revealed a substantial amount of viral gene expression heterogeneity, although the distributions differed somewhat from what we observed with Cal07 (**Fig. S2**). These differences may be partially due to an artifact of the QC process by which we eliminated cell doublets from our analysis (see Methods and **Fig. S4** for details). Regardless, Perth09-infected cells exhibited the same pattern of discordance between NS and the rest of the genome segments that we observed with Cal07 (**Fig. 6C, S3**).

To identify broader patterns of viral gene expression, we performed K-means clustering based on expression of the eight viral gene segments for all Cal07- and Perth09-infected cells that were positive for all viral transcripts. Clustering patterns were highly similar for both viruses and revealed the existence of three distinct infected cell populations that largely subdivided cells based on both total viral gene expression levels and NS segment expression (**Fig. 6D**). Thus, human H1N1 and H3N2 viruses share a common pattern of single cell viral gene expression in which NS segment expression diverges from that of the rest of the viral genome in a subset of infected cells.

### Single cell variation in NS segment expression is a major driver of the IFN response to Perth09, but not Cal07 infection

To determine whether these distinct clusters of viral gene expression were associated with distinct patterns of host transcription, we compared host transcript expression of infected cells in each cluster to the other two clusters using both MAST and NBID. This approach identified dozens of host genes differentially expressed in each cluster (**Table S11-16**). For Cal07, no IFN genes were significantly differentially expressed between clusters, but for Perth09 there were clear differences in the expression of numerous host antiviral defense genes between the three clusters. The expression frequencies and/or levels for type III IFNs (IFNL1 and IFNL2) and selected ISGs (IFIT2, ZC3HAV1) were significantly higher in cluster 2, the cluster with low NS expression, compared with the other clusters (**Fig. 6E-H**). Thus, for Perth09 but not Cal07, single cell heterogeneity in NS segment expression level is a major determinant of the activation of the innate antiviral response.

### H1N1 and H3N2 strains can differ significantly in single cell patterns of IFN and ISG transcription

Finally, we compared the activation of host innate anti-viral gene expression at the single cell level between Cal07 and Perth09. Mirroring previous observations, expression of type I and III IFNs were only observed in a tiny fraction of Cal07-infected cells (**Fig. 7A**) (13,39). For type I IFN (but probably not type III IFN), this could partially be a function of the relatively late timepoint that we examined, as IFNbeta expression typically peaks earlier during infection (40). Expression of IFNalpha2 was also rare in bystander cells (**Fig. 7B**). We also observed widespread expression of multiple ISGs within Cal07-infected cells indicating the successful activation of host antiviral defenses within infected cells despite the low frequency of IFN expression, as has been described previously (11,41). Similar to what we observed with IFNalpha2, ISG expression was minimal in bystander cells, suggesting that the inhibition of IFN induction by Cal07 is sufficient to largely prevent paracrine ISG activation in this host system (**Fig. 7C**).

**Fig. 7.**
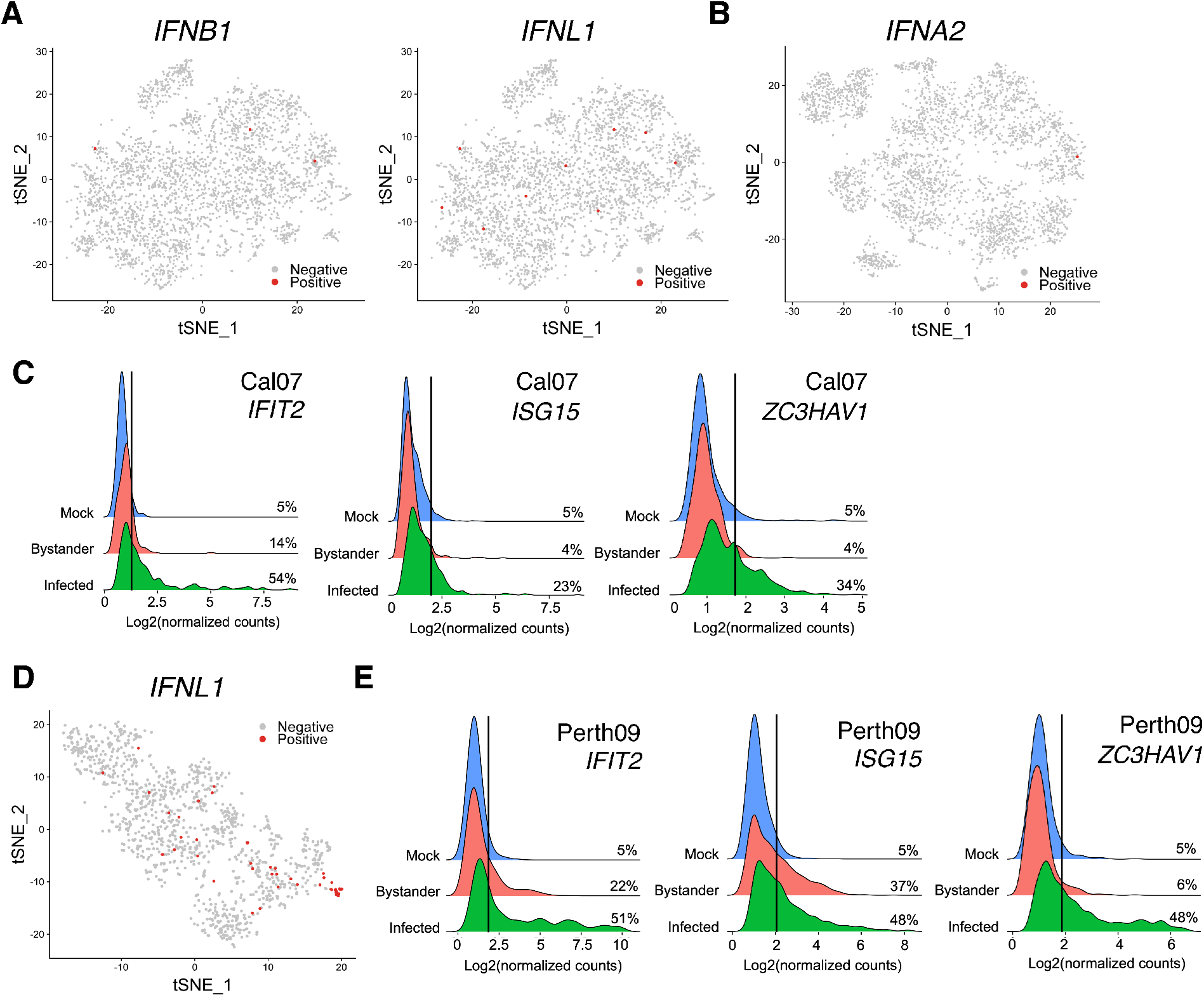
H1N1 and H3N2 strains differ significantly in single cell patterns of IFN and ISG transcription. **(A)** tSNE plots of all Cal07-infected A549 cells colored based on whether IFNB1 and IFNL1 are detected. Results are representative of all type III IFNs examined. **(B)** tSNE plot of all bystander A549 cells colored based on whether IFNA2 is detected. Similar results are found for other IFN-α subtypes. **(C)** Histograms comparing distributions of per-cell normalized counts for the indicated host transcripts across the three libraries (mock, bystander, and infected) during Cal07 infection. Vertical lines indicate 95% cutoff thresholds based on mock libraries, with percentages of cells in each library above the thresholds shown on right. **(D)** tSNE plot of all Perth09-infected A549 cells colored based on whether IFNL1 is detected. Results are representative of all type III IFNs examined. **(E)** Same as in (C) for Perth09 infection.

During Perth09 infection, type III IFN transcription was roughly 20-fold more frequent compared with Cal07, indicating a significant difference in the ability of the host to initiate an IFN response to infection by these two strains (**Fig. 7D**). Consistent with this, some (but not all) ISGs were expressed more frequently in bystander cells during Perth09 infection compared with Cal07, suggesting that Perth09 is less able to prevent paracrine activation of bystander cells (**Fig. 7E**). Altogether, these data reveal clear differences in single cell patterns of IFN and ISG transcription between the human H1N1 and H3N2 strains tested.

## DISCUSSION

The host response to infection arises from the combined responses of many individual cells, both infected and uninfected. To understand the factors that govern host responses at the tissue and organismal levels, it is critical to define patterns of variation in single cell infection responses. Here, we demonstrate that IAV infection gives rise to a heterogeneous collection of divergent transcriptional responses at the single cell level. Notably, this heterogeneity occurred within a single cell type, under conditions where per-cell viral input and infection timing were normalized. Single cell responses *in vivo* may be significantly more variable. By examining patterns of viral gene expression heterogeneity at the single cell level, we uncovered a surprising dynamic where NS segment expression diverges from that of the other genome segments. This pattern was shared between the human seasonal strains that we tested and played a major role in regulating the host antiviral response to H3N2 infection. These data establish a clear role for viral heterogeneity in modulating the host response to infection and highlight the power of single cell approaches to reveal new determinants of viral infection outcomes.

Variation in gene expression patterns between individual infected cells appears to arise from a combination of (1) genetic and genomic variation between individual virions, (2) underlying heterogeneity within host cells, and (3) stochastic variation in infection processes. In line with previous reports of significant genetic and phenotypic heterogeneity within A459 and other transformed cell line populations, we observed substantial transcriptional heterogeneity between uninfected A549 cells that could not be simply explained by cell cycle status (42–44). As a result, virions that enter different cells establish infection within differing cellular environments that may be more or less supportive and primed for differing transcriptional responses to infection. This effect is likely enhanced by the stochastic variation inherent in different viral life cycle stages (45).

Layered on top of this intrinsic cellular heterogeneity and stochasticity is the genetic and genomic heterogeneity characteristic of IAV populations (5). We and others have previously demonstrated that well-characterized viral modulators of host cell function such as NS1 and PA-X are not ubiquitously expressed during infection (6,12). Here, we show that this variability in viral gene expression between individual infected cells has clear consequences for the host cell transcriptional response. Interestingly, we found that two levels of viral gene expression heterogeneity influence single cell infection outcomes: variation in the presence or absence of a given viral gene and, for cells where a gene is expressed, variation in expression level. This effect was most pronounced for the NS segment where variation in expression levels between cells was associated with significant differences in the host transcriptional response to infection. Altogether, it is clear that future efforts to understand the role of critical viral proteins such as NS1 in shaping infection outcome will have to account for heterogeneity in single cell expression patterns.

The comparison of individual viral gene segment expression across thousands of infected cells revealed surprisingly complex relationships between the NS segment and the rest of the genome. Within a subset of infected cells, NS expression was markedly decreased compared to the other viral genes. Importantly, this dynamic was not universal, as NS expression was well-correlated with expression of the other gene segments in some clusters of infected cells. This suggests that multiple distinct states of viral gene expression can arise during infection. Similar to what has been shown during genome packaging, our data suggest that specific interactions between gene segments play an important role in regulating overall viral gene expression patterns (46–48). Future efforts will aim to understand the mechanism behind the unique expression pattern of the NS segment.

The comparison of Cal07 and Perth09 revealed clear commonalities and differences in single cell expression patterns between the two viruses that may have relevance for understanding differences in the apparent pathogenicity of different human H1N1 and H3N2 viruses. The most obvious difference between the two viruses was seen in the activation of IFN and ISG transcription. In addition, while NS segment expression status significantly influenced the host response to both viruses, only Perth09 showed a clear relationship between NS levels and the activation of the host antiviral machinery. This finding is in line with previous work that showed that the 2009 pandemic H1N1 NS1 protein was less effective at suppressing transcription of human IFNs and ISGs compared with NS1 genes from other human IAV isolates (49). While *de novo* NS segment transcription appeared to be largely dispensable for blocking IFN and ISG upregulation by Cal07, overall expression frequencies for select IFNs and ISGs were actually lower during Cal07 infection compared with Perth09, indicating that Cal07 is more effective at suppressing IFN activation in A549 cells. Overall, our data make clear that there is still a lot that we do not understand about the interplay between viral gene expression patterns, viral genotype, and the host antiviral response.

In this study, we took steps to eliminate two other sources of viral heterogeneity that are likely common during natural IAV infection. The first is presence of defective interfering particles (DIPs), which we minimized by generating our virus stocks under low MOI conditions. DIPs appear to be common within IAV populations, even in humans, and can have complicated effects on both viral and host gene expression (13,50–55). We would predict that the presence of DIPs would further increase overall heterogeneity between individual cell responses to the virus. It is worth noting that we could not reliably detect the presence of DIP-associated transcripts within our data due to the unavoidably incomplete coverage of the viral gene segments generated by our short-read sequencing approach. The second likely source of additional heterogeneity that we excluded from this study is variability in the cellular MOI. We and others have demonstrated that cellular co-infection can be common *in vivo* (7,56–58). This suggests that the number of viral genomes entering individual cells is likely quite variable. We recently showed that this variability in cellular MOI can have distinct phenotypic consequences, both for viral replication dynamics and for IFN induction (21). Altogether, it appears likely that the heterogeneity that we describe here underrepresents what would be observed *in vivo*.

Our results extend the previous results of other groups including our own in establishing the enormous amount of heterogeneity in viral gene expression that occurs at the single cell level, even under experimental conditions designed to minimizes sources of variability (6,10,12,45,59). Critical phenotypes such as viral load dynamics, transmissibility, and pathogenicity must emerge from the collective output of heterogeneous populations of infected cells. This raises questions of how selection may act upon patterns of viral heterogeneity to alter these emergent phenotypes and the extent to which these heterogeneity patterns are under viral genetic control. These questions are especially pertinent for segmented viruses like IAV but are likely relevant across diverse virus families.

Altogether, our results help establish the importance of considering the roles of viral and host cell heterogeneity in influencing the pathogenesis of viral infections. Similar to the way that viral populations are now viewed, our data clearly establishes that the host response to infection should be seen as a heterogeneous assemblage of single cell responses that collectively give rise to the bulk phenotypes that are generally measured. This creates the potential for complex interactions between responding cell subsets and raises the question of how such a heterogeneous system is effectively regulated. It also raises the possibility that viral and host response dynamics may be disproportionately driven by small subsets of cells that are obscured during bulk analysis. Dissection of these diverse constituents is likely to reveal new mechanisms that govern the pathogenesis of influenza virus infection.

## METHODS

### Plasmids

The A/California/04/09 and A/Perth/16/2009 reverse genetics plasmids were generous gifts from Drs. Jonathan Yewdell and Seema Lakdawala, respectively. Plasmids encoding A/California/07/09 were generated by introducing A660G and A335G substitutions into HA and NP respectively, to match the amino acid sequences of A/California/07/09 HA and NP (NCBI accession numbers CY121680 and CY121683).

### Cells

Madin-Darby canine kidney (MDCK) and human embryonic kidney HEK293T (293T) cells were maintained in Gibco’s minimal essential medium with GlutaMax (Life Technologies). Human lung epithelial A549 cells were maintained in Gibco’s F-12 medium (Life Technologies). MDCK and A549 cells were obtained from Dr. Jonathan Yewdell; 293T cells were obtained from Dr. Joanna Shisler. All media were supplemented with 8.3% fetal bovine serum (Seradigm). Cells were grown at 37°C and 5% CO2.

### Viruses

Recombinant A/California/07/09 (Cal07) and A/Perth/16/2009 (Perth09) viruses were rescued via the 8-plasmid reverse genetics approach. For the rescue of both viruses, sub-confluent 293T cells were co-transfected with 500ng of the following plasmids: pDZ::PB2, pDZ::PB1, pDZ::PA, pDZ::HA, pDZ::NP, pDZ::NA, pDZ::M, and pDZ::NS, using JetPrime (Polyplus) according to the manufacturer’s instructions. Plaque isolates derived from rescue supernatants were amplified into seed stocks in MDCK cells. Working stocks were generated by infecting MDCK cells at an MOI of 0.0001 TCID_50_/cell with seed stock and collecting and clarifying supernatants at 48 hpi. All viral growth was carried out in MEM with 1 μg/ml trypsin treated with L-(tosylamido-2-phenyl) ethyl chloromethyl ketone (TPCK-treated trypsin; Worthington), 1 mM HEPES, and 100 μg/ml gentamicin. The titers of the virus stocks were determined via standard 50% tissue culture infectious dose (TCID_50_) assay.

### Viral infection and cell sorting for single cell RNAseq

Confluent A549 cell monolayers in 3 T-25 flasks were infected with Cal07 (or Perth09) at MOI of 0.01 TCID_50_/cell for 1 h. At 1 hpi, monolayers were washed with phosphate-buffered saline (PBS) and incubated in serum-containing F-12 medium. At 3 hpi, the medium was changed to MEM with 50 mM HEPES and 20 mM NH4Cl to block viral spread. At 16 hpi, monolayers were trypsinized and combined into single-cell suspension and washed with PBS. Cal07-infected cells were stained with Alexa Fluor 488-conjugated mouse anti-HA monoclonal antibody (mAb) EM4-CO4 (gift from Dr. Patrick Wilson) and Alexa Fluor 647-conjugated mouse anti-M2 mAb O19 (gift from Dr. Jonathan Yewdell). Perth09-infected cells were first stained with human anti-HA stem antibody FI6 (Gift from Dr. Adrian McDermott) and Alexa Fluor 647-conjugated mouse anti-M2 mAb O19, and then stained with Alexa Fluor 488-conjugated donkey anti-human IgG (Jackson ImmunoResearch). After staining, cells were washed with PBS twice, and single live cells were sorted as “infected” or “bystander” populations based on the expression of HA and M2 on a BD FACS ARIA II sorter. Importantly, uninfected A549 cells from a separate flask were also trypsinized, stained, and sorted as “mock” population which served as a negative control.

### Single cell RNAseq cDNA library generation

Sorted cell samples were counted and checked for viability on a BD20 cell counter (BIO-RAD) before they were diluted to equivalent concentrations with an intended capture of 4000 cells/sample. Each individual sample was used to generate individually barcoded cDNA libraries using the 10x Chromium Single Cell 3’ platform (Pleasanton, CA) following the manufacturer’s protocol. The Chromium instrument separates single cells into Gel Bead Emulsions (GEMs) that facilitate the addition of cell-specific barcodes to all cDNAs generated during oligo-dT-primed reverse transcription. The experiment with Cal07 used V2 reagent and the experiment with Perth09 used V3 reagent.

### Illumina Library preparation and sequencing

Following ds-cDNA synthesis, individually barcoded libraries compatible with the Illumina chemistry were constructed. The libraries were sequenced on an Illumina NovaSeq 6000 using S4 flowcell for the experiment with Cal07 and S3 flowcell for the experiment with Perth09.

### Single cell RNAseq analysis

The three Cal07-associated 10x Chromium Single Cell v2 libraries (infected, bystander, and mock) were demultiplexed and reference mapped using Cell Ranger Count (version 2.2) for alignment to a combined human+virus reference (human: hg38, version 1.2.0; Cal07: Genbank Accessions: CY121680-CY121687), and then combined using Cell Ranger Aggr. The three Perth09-associated 10x Chromium Single Cell v3 libraries (infected, bystander, and mock) were processed and combined using Cell Ranger (version 3.1) for alignment to a combined human+virus reference (human: hg38, version 1.2.0; Perth09: Genbank Accessions: KJ609203.1-KJ609210.1). The resulting raw count matrix for each virus set was imported into an R pipeline using SimpleSingleCell (60), where it was filtered for empty droplets (61), low feature cells (i.e. droplets removed if < 400 features/cell), low expressing features (i.e. droplets removed if < 4cells/feature), and potential doublets (droplets removed if UMI counts of host transcripts > 2-fold of the median host UMI count number).

During the QC process for the Perth09 infection experiment, roughly half of the infected cells registered as “doublets” based on total host mRNA content, despite our filtering out cell doublets during flow sorting (**Fig. S4**). This appears to be a Perth09-specific effect, as we did not see this during Cal07 infection. While it is possible that many of these are actually single cells in which total RNA amounts are elevated due to infection, we removed these cells from our analysis.

The filtered matrix was scale normalized using the deconvolution method described and implemented in the scran R package (60). Overall cellular infection status and individual virus gene presence/absence was determined to be above background by comparison to expected background counts generated from the subset of cells from the “Bystander” library containing virus counts (assumed to be background molecules). For bystander cells with virus-mapping reads, we calculated the 95th percentile values for total virus molecule cellular proportions (‘Total Cellular Virus’) as well as for each of the individual gene segments (‘Total Cellular Virus Gene’) (**Fig. S5**) and used these values to calculate the ‘Expected Virus Background Proportion’ for all cells. These values represent the threshold below which we assume viral signal is due to background cross-contamination rather than true viral gene expression. Cell cycle status was determined by running the Cyclone tool in the scran R package (60). The filtered, normalized, and annotated matrix was then imported into a Seurat pipeline for additional analysis and visualization (62), including differential gene expression analysis using the MAST (33) and NBID tools (32), graph-based clustering of the cells (63), and PCA/tSNE dimensional reduction visualization.

Differential Gene Analysis (DGE) gene lists were produced from the filtered, normalized count matrices using R packages MAST and NBID (32,33). Individual MAST and NBID results were intersected using a custom Perl script to obtain final lists. DGE lists for missing virus genes were generated by first sub-setting the count matrices to contain only ‘Infected’ status cells and then using the individual virus gene status factors to test for cells with ‘Present’ versus ‘Absent’ status. DGE lists for individual cell clusters were generated by first sub-setting for ‘Infected’ status cells and then by non-zero cluster ID status. Tests were then performed for each cluster versus all other clusters.

All code used for single cell analysis, along with associated documentation, is available from: https://github.com/BROOKELAB/SingleCell. All sequencing data will be available from the NCBI GEO at the time of publication.

### Multivariate homogeneity of groups dispersions analysis

In the analyses, each host gene represents a variable/coordinate and thus each cell can be seen as a point in the multivariate space. Multivariate homogeneity of groups dispersion analysis (30) was performed on log transformed host gene expressions of the two group of cells, i.e. mock cells and infected cells, using the betadisper function in the R package (http://www.R-project.org/). Distances between the points (i.e. cells) and their respective group centroid in the principal coordinates were then used to test homogeneity of variances and calculate the p-value. Note that similar analyses were performed on linear host gene expression values and results remain the same, i.e. overall host gene expressions are significantly more heterogenous than those in mock cells.

### K-means clustering

The clusters are identified by applying standard k-means clustering analysis to partition the joint 8-viral gene segment expressions (on a log scale) into three distinct clusters (k=3).

## ACKNOWLEDGEMENTS

We are grateful to Dr. Alvaro Hernandez and Ms. Chris Wright of the DNA Services Lab within the Roy J. Carver Biotechnology Center for expert assistance in experimental planning and the preparation and sequencing of single cell RNAseq libraries. We also would like to thank Dr. Chris Fields of HPCBio, also within the Roy J. Carver Biotechnology Center, for assistance in troubleshooting analysis. This work has been generously funded by NIAID (1R01AI139246-01A1), the DARPA INTERCEPT program (DARPA INTERCEPT W911NF-17-2-0034 and R-00676-19-0), and the Roy J. Carver Charitable Trust (17-4905).

## SUPPLEMENTAL FIGURES

**Figure S1. All pairwise Cal07 viral gene correlation plots.** Normalized per cell copy numbers for the indicated Cal07 genes plotted against each other. Data only show infected cells that are positive for all viral gene segments.

**Figure S2. Single cell heterogeneity in viral gene expression during low MOI Perth09 infection. (A)** Distribution of Perth09-infected A549 cells, binned by the fraction of total cellular poly(A) RNA that is viral in origin. **(B)** Fraction of total poly(A) RNA per cell that maps to the indicated viral gene segment. Each dot represents a single cell, cells with no detectable reads mapping to the indicated segment are arbitrarily assigned a value of 0.000007 to show up on the log10 scale. **(C)** Fraction of all Perth09-infected cells that have detectable levels of transcripts derived from the indicated viral gene segments. **(D)** All Perth09-infected cells binned by the total number of detectable viral gene segments, with the actual numbers of cells in each group detailed above.

**Figure S3. All pairwise Perth09 viral gene correlation plots.** Normalized per cell copy numbers for the indicated Perth09 genes plotted against each other. Data only show infected cells that are positive for all viral gene segments.

**Figure S4. Distributions of single cell host mRNA counts and doublet calling for Cal07 and Perth09 infection.** Single cell distributions of host UMI counts for mock, bystander, and infected cell libraries during **(A)** Cal07 and **(B)** Perth09 infection. In each group, the vertical line indicates a value equal to the median UMI count of host transcripts in combined mock, bystander, and infected cells, multiplied by 2. All cells with host UMI counts greater than this value are considered doublets and are excluded from subsequent analyses.

**Figure S5. Determination of cutoff thresholds for infection status of cells and expression status of viral gene segments.** Single cell distributions of percentages of total mRNA in bystander cells during Cal07 infection that is either **(A)** all viral mRNA, or **(B)**PB1-derived mRNA. The vertical line indicates the cutoff threshold equal to the 95^th^ percentile value in each plot. All cells with total viral mRNA fraction greater than the cutoff in (A) are considered infected, and all cells with PB1 mRNA fraction greater than the cutoff in (B) are considered positive for PB1. This cutoff calling method is used for each of the eight viral gene segments for both Cal07 and Perth09.

## SUPPLEMENTAL TABLES

**Table S1: All host genes differentially expressed between Seurat clusters of Cal07 infected cells**

**Table S2: Combined DGE list based on Cal07 PB2 expression status within infected cells**

**Table S3: Combined DGE list based on Cal07 PB1 expression status within infected cells**

**Table S4: Combined DGE list based on Cal07 PA expression status within infected cells**

**Table S5: Combined DGE list based on Cal07 HA expression status within infected cells**

**Table S6: Combined DGE list based on Cal07 NP expression status within infected cells**

**Table S7: Combined DGE list based on Cal07 NA expression status within infected cells**

**Table S8: Combined DGE list based on Cal07 M expression status within infected cells**

**Table S9: Combined DGE list based on Cal07 NS expression status within infected cells**

**Table S10: Combined DGE list showing all host genes differentially expressed within infected cells based on expression status of individual viral gene segments**

**Table S11: Combined DGE list showing host genes differentially expressed in cluster 1 (from K means clustering of all Cal07 infected cells positive for all viral genes)**

**Table S12: Combined DGE list showing host genes differentially expressed in cluster 2 (from K means clustering of all Cal07 infected cells positive for all viral genes)**

**Table S13: Combined DGE list showing host genes differentially expressed in cluster 3 (from K means clustering of all Cal07 infected cells positive for all viral genes)**

**Table S14: Combined DGE list showing host genes differentially expressed in cluster 1 (from K means clustering of all Perth09 infected cells positive for all viral genes)**

**Table S15: Combined DGE list showing host genes differentially expressed in cluster 2 (from K means clustering of all Perth09 infected cells positive for all viral genes)**

**Table S16: Combined DGE list showing host genes differentially expressed in cluster 3 (from K means clustering of all Perth09 infected cells positive for all viral genes)**

