## Supplemental figures for "A common pattern of influenza A virus single cell gene expression heterogeneity governs the innate antiviral response to infection"

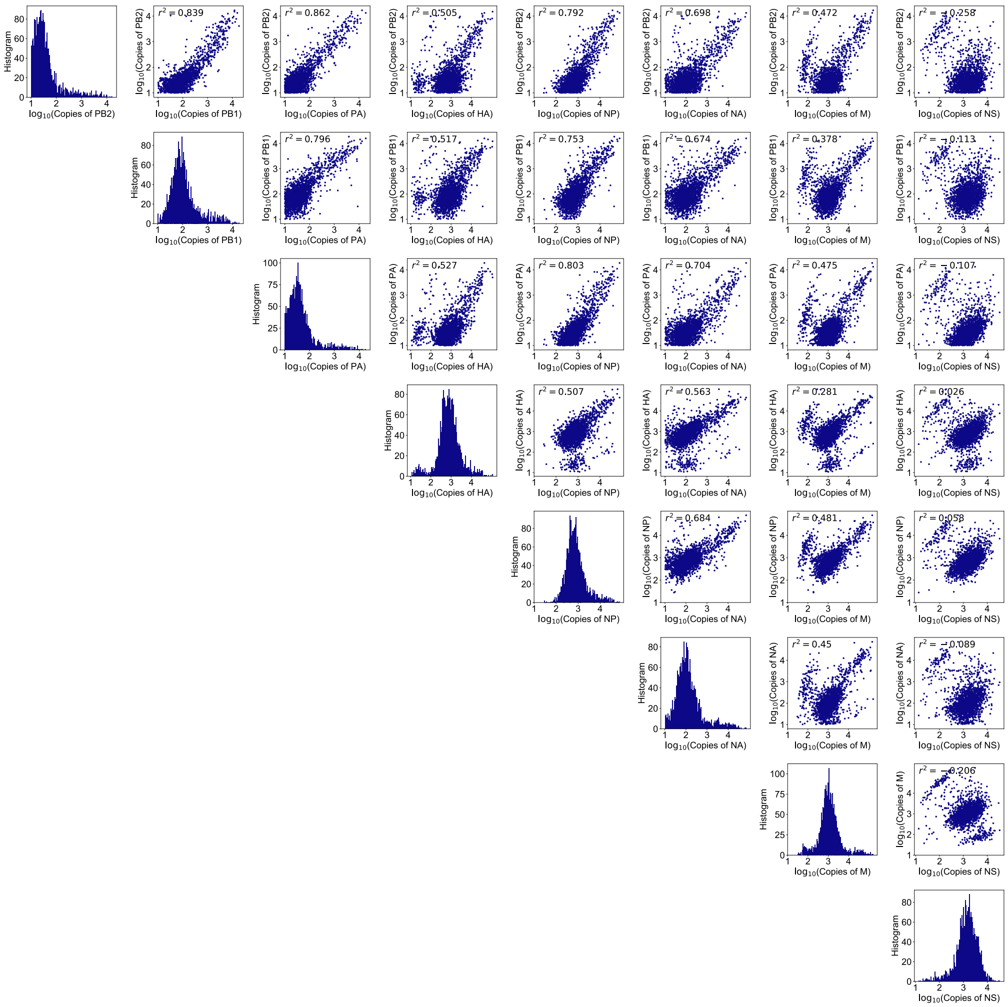

**Figure S1. All pairwise Cal07 viral gene correlation plots.** Normalized per cell copy numbers for the indicated Cal07 genes plotted against each other. Data only show infected cells that are positive for all viral gene segments.

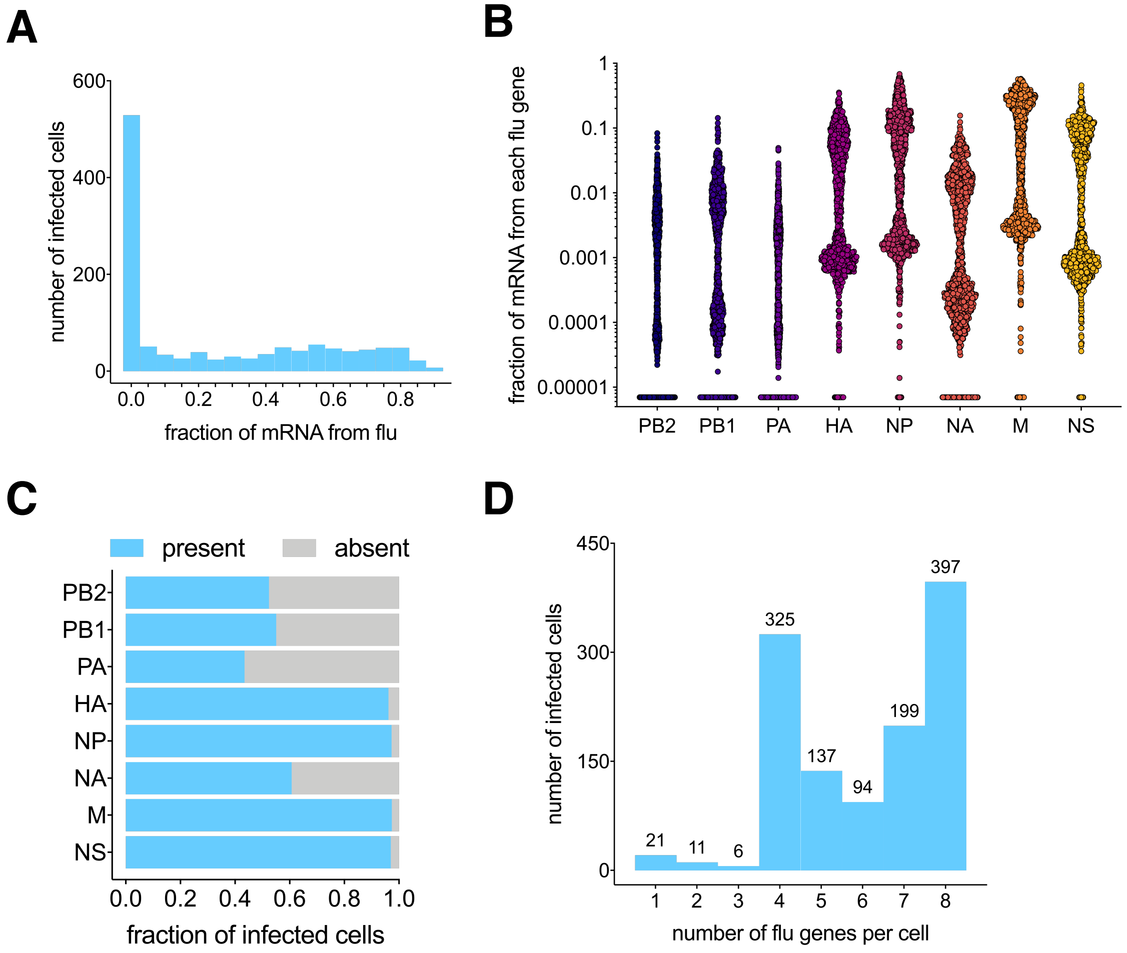


**Figure S2. Single cell heterogeneity in viral gene expression during low MOI Perth09 infection. (A)** Distribution of Perth09-infected A549 cells, binned by the fraction of total cellular poly(A) RNA that is viral in origin. **(B)** Fraction of total poly(A) RNA per cell that maps to the indicated viral gene segment. Each dot represents a single cell, cells with no detectable reads mapping to the indicated segment are arbitrarily assigned a value of 0.000007 to show up on the log10 scale. **(C)** Fraction of all Perth09-infected cells that have detectable levels of transcripts derived from the indicated viral gene segments. **(D)** All Perth09-infected cells binned by the total number of detectable viral gene segments, with the actual numbers of cells in each group detailed above.

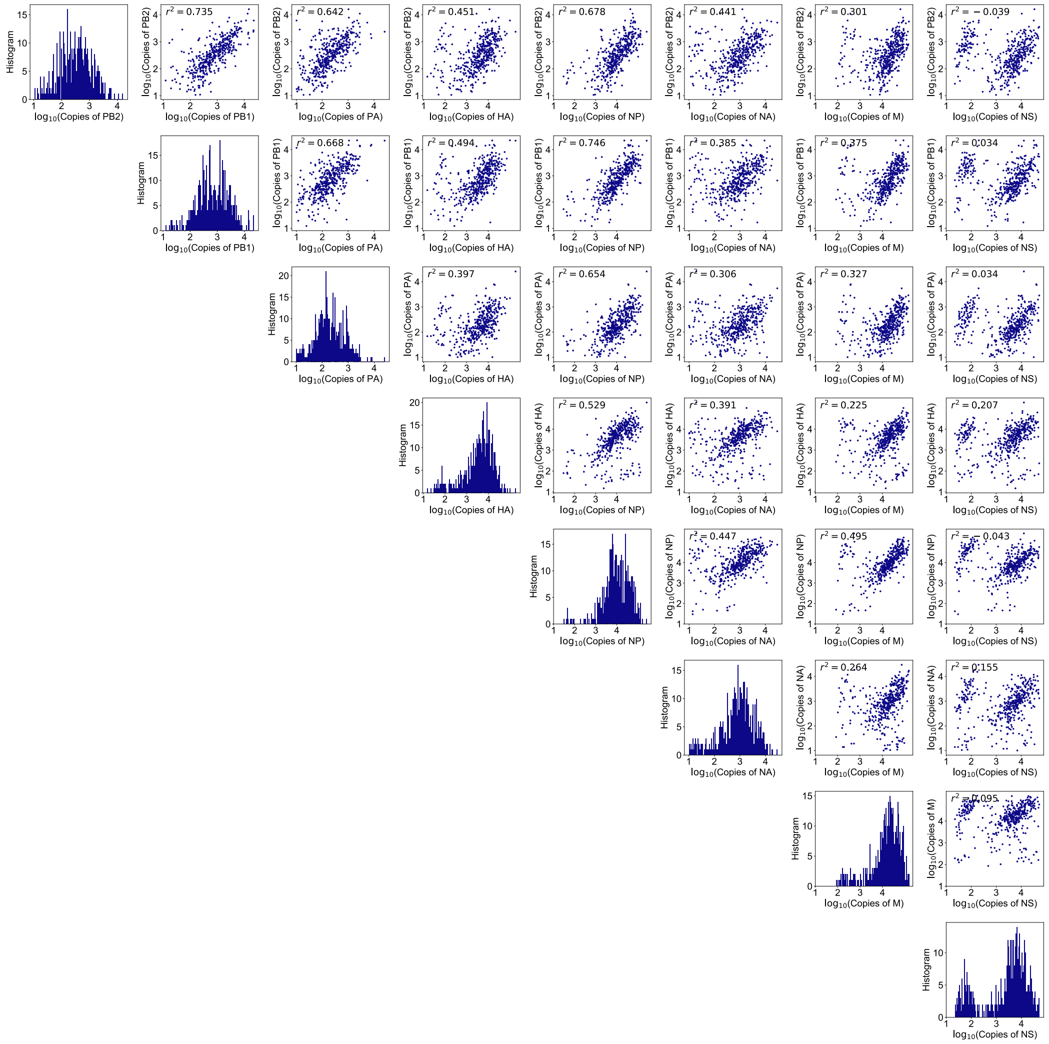


**Figure S3. All pairwise Perth09 viral gene correlation plots.** Normalized per cell copy numbers for the indicated Perth09 genes plotted against each other. Data only show infected cells that are positive for all viral gene segments.

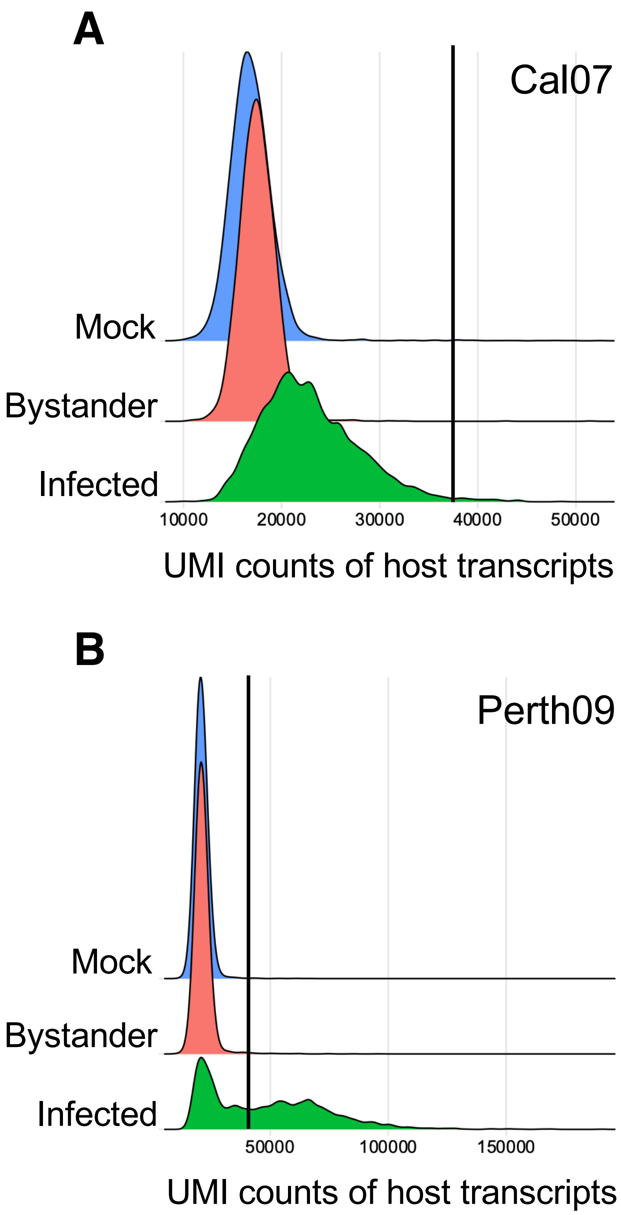


**Figure S4. Distributions of single cell host mRNA counts and doublet calling for Cal07 and Perth09 infection.** Single cell distributions of host UMI counts for mock, bystander, and infected cell libraries during **(A)** Cal07 and **(B)** Perth09 infection. In each group, the vertical line indicates a value equal to the median UMI count of host transcripts in combined mock, bystander, and infected cells, multiplied by 2. All cells with host UMI counts greater than this value are considered doublets and are excluded from subsequent analyses.

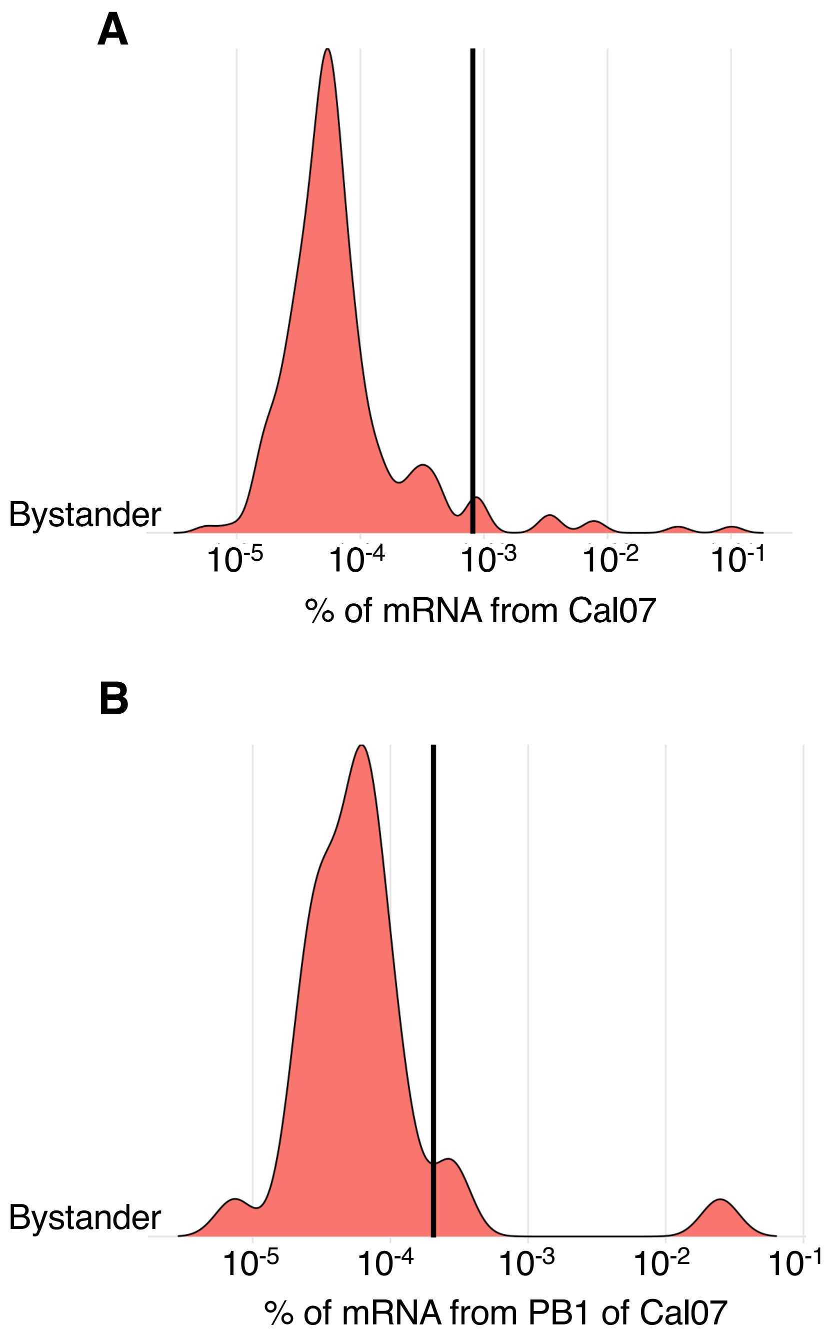


**Figure S5. Determination of cutoff thresholds for infection status of cells and expression status of viral gene segments.** Single cell distributions of percentages of total mRNA in bystander cells during Cal07 infection that is either **(A)** all viral mRNA, or **(B)** PB1-derived mRNA. The vertical line indicates the cutoff threshold equal to the 95th percentile value in each plot. All cells with total viral mRNA fraction greater than the cutoff in (A) are considered infected, and all cells with PB1 mRNA fraction greater than the cutoff in (B) are considered positive for PB1. This cutoff calling method is used for each of the eight viral gene segments for both Cal07 and Perth09.
